# An analysis and optimization of growth condition requirements of the fast-growing bacterium *Vibrio natriegens*

**DOI:** 10.1101/775437

**Authors:** Celine Örencik, Sara Müller, Thomas Kirner, Egon Amann

## Abstract

The fast-growing Gram-negative bacterium *Vibrio natriegens* is an attractive host for a range of applications in molecular biology and biotechnology. Moreover, the remarkable speed of growth of *Vibrio natriegens* poses fundamental questions on bacterial physiology and metabolism, energy production, DNA replication and protein synthesis, besides others. In order to address such questions, a solid understanding of the physiological and physical/chemical basis of growth requirements is essential. Here we report a systematic analysis of i) various growth media composition, ii) incubation temperature, iii) pH dependence, and iv) salt concentration requirements for optimal growth of *V. natriegens* strain DSMZ 759. As a result of the studies, the following optimal conditions were Established: LB medium with 2.5 % NaCl, pH 7.0 – 8.5 and incubation at 37°C under aerobic conditions. Incubation temperatures above 37 °C slows growth significantly. Incubation temperatures below 37 °C slows growth, but at a lower rate. Incubation at or below 28 °C should be avoided. Under such optimized, standard laboratory conditions, a doubling time of t_d_ = 13.6 minutes was observed for *V. natriegens* measured in mid-log growth phase. The optimized conditions presented here for the growth of *V. natriegens* can be easily applied in any standardly equipped laboratory. For comparison, identical growth conditions for *Escherichia coli* were analyzed and are presented as well.

**IMPORTANCE:** Goal of this study was to understand the physiological growths requirements of *V. natriegens* in routine microbiology and molecular biology laboratory settings. The result is a standardized protocol for the optimized growth of the naturally isolated (wild type) *V. natriegens* strain DSMZ 759. This protocol can be employed for routine application of *V. natriegens* for any kind of biochemical, molecular biology and genomic studies and utilization under normal laboratory conditions used by many routinely equipped laboratories.

## INTRODUCTION

The prototroph, gram-negative, non-pathogenic marine bacterium *Vibrio natriegens* was originally isolated from salt marsh muds of Sapelo Island, GA, and described by Payne et al. in 1961 (1). Initially, this bacterium was taxonomically classified into the genus *Pseudomonas* (*P. natriegens*), but later redescribed as *Beneckea natriegens.* Finally it was recognized as a member of the genus *Vibrio* by Austin et al. in 1978 (2). In 1962 Eagon firstly described the unusually short generation time of *Vibrio natriegens* of 9.8 minutes applying incubation at 37 °C in brain heart infusion (BHI) with vigorous shaking (3). Nowadays, 9.8 minutes is still the shortest doubling time ever observed in biological investigations, and interesting speculations can be made about the limits of growth rates in prokaryotic microorganisms (4). However, even shorter doubling times have experimentally been achieved using a small (50 mL) glass fermenter under optimized conditions such as temperature, nutrients, and air supply (5, 6). The fastest growth with doubling times below 7 min was obtained only when the optical density (OD) of liquid cultures was very low (OD at 600 nm of between 0.05 and 0.1) and the cultures were under strong aeration (4).

In a different study, a doubling time of 9.4 minutes has recently been reported under optimized fermentation conditions using BHIN complex medium as well as in minimal medium supplemented with various industrially relevant substrates (7). This study used *V. natriegens* as a production host for industrial biotechnology applications and showed that the observed growth rate is at least two times higher than those of traditionally employed microbial systems such as *Escherichia coli, Bacillus subtilis, Corynebacterium glutamicum* and yeast.

The draft genome sequence of *V. natriegens* strain DSMZ 759 was recently published (4). The *V. natriegens* genome consists of two chromosomes of a combined size of 5,200,362 bp with approximately 4,788 coding sequences, including 3,542 with a putative functional annotation, 14 rRNA-encoding genes, and 71 tRNA-encoding genes (4). In a different study using a different *V. natriegens* strain, the first complete *V. natriegens* genome, consisting of two chromosomes of 3,248,023 bp and 1,927,310 bp that together encode 4,578 open reading frames, was published (8).

*V. natriegens’* high potential for protein synthesis results from an increase in ribosome numbers with increasing growth rates and from a large number of rRNA operons in addition to extremely strong rRNA promoter activity (9). Based on these features, cell-free expression systems employing *V. natriegens* compounds have been established (10, 11, 12).

Intrigued by this extremely fast growth dynamics, *V. natriegens* was proposed and is already applied as a “new” host for applications in molecular biology, biotechnology and genomics and is on its way to replace *E. coli*, at least in part, for such applications (4, 8, 13).

Here we report a study comparing *V. natriegens*’ with *E. coli’s* growth requirements (growth media, pH, salinity, and temperature) with the aim to establish a simple and reliable protocol for the standard, routine laboratory without the quest for “fastest” growth records. The resulting protocol should be useful for a variety of applications of *V. natriegens* in molecular biology, microbiology and industrial production of heterologous proteins as well as studying its metabolism in further detail.

## RESULTS

### Medium Optimization

We compared *V. natriegens* growth characteristics in LB medium, nutrient agar and M9 minimal agar. For comparison, *E. coli* grown in LB medium with 1.5 % NaCl was included in the experiment (Fig. 1A). Best growth dynamics were observed with LB medium with 2.5 % and 1.5 % NaCl. However, growth dynamics of nutrient agar supplemented with 2.5 % NaCl were comparably good. Nutrient agar supplemented with 1.5 % NaCl resulted in delayed bacterial growth. In M9 minimal agar not supplemented with NaCl growth was slow and decayed after 4 hours. *E. coli* grown in LB medium with a NaCl concentration of 1.5 % reached after 16 hours and after a long lag phase also a high final OD, although lower when compared with *V. natriegens* maximal reach. Growth curves of *E. coli* cultures under the same medium conditions were also recorded and are shown in Fig. S4.

**FIG 1.**
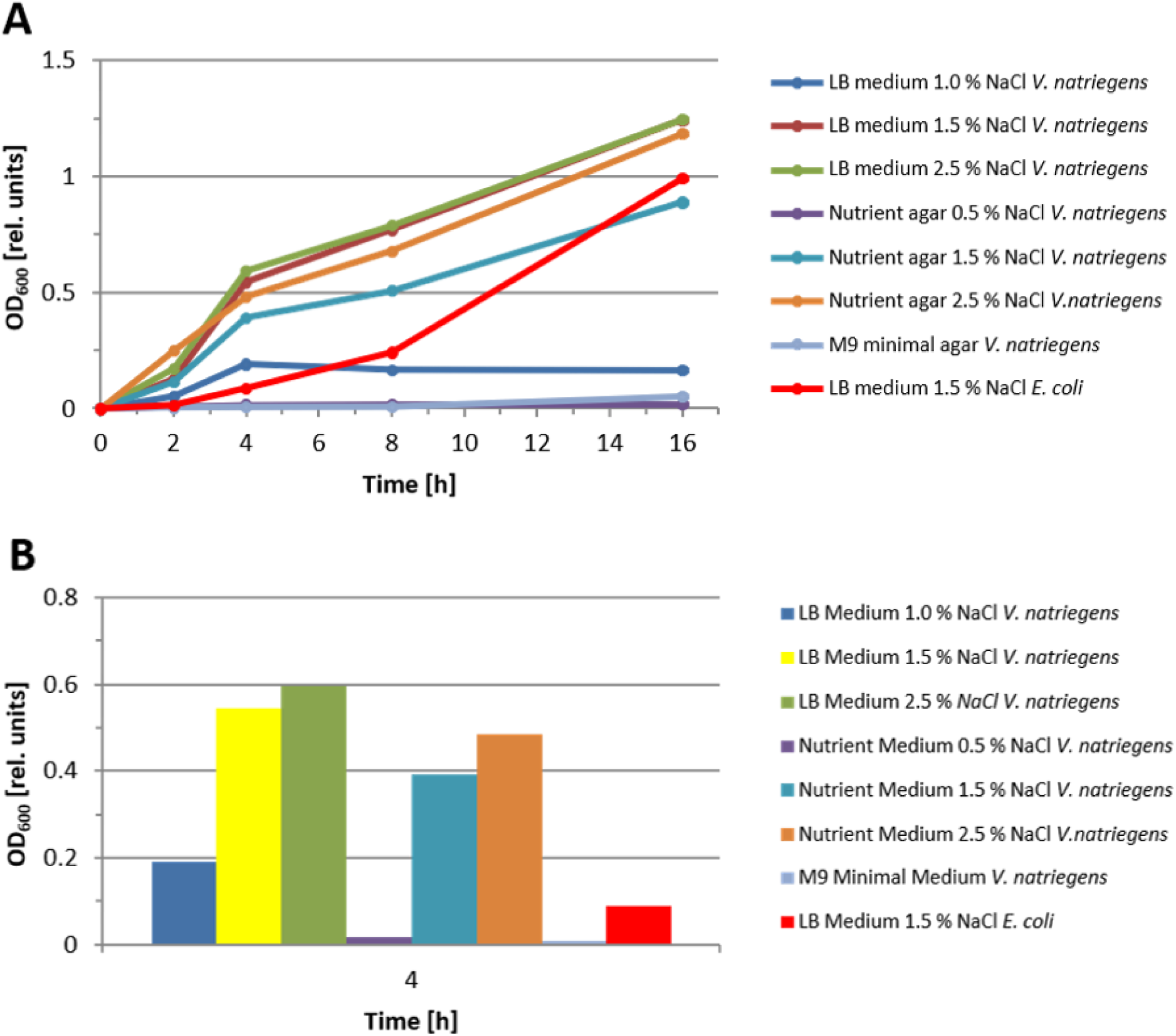
Growth of *V. natriegens* in different standard media over time (A) and shown after 4 hours growth as single time point (B). For comparison, growth of *E.coli* is shown in red. The respective medium composition is given in the Materials and Methods section.

Based on this result we conclude that LB medium with a NaCl concentration of 2.5 % to be the best medium of the three media tested to support good growth of *V. natriegens*. For better comparison of the tested media and salt concentrations, the 4 hours ODs are depicted in a block diagram (Fig. 1B).

### Temperature Optimization

We wanted to investigate the optimal temperature for efficient growth of *V. natriegens.* LB medium cultures supplemented with 2.5 % NaCl were inoculated with the same starting amounts of bacteria and incubated at 28 °C, 31 °C, 34 °C, 37 °C and 40 °C, respectively. Samples were taken at the indicated times and the OD was recorded. *E. coli* grown under identical medium composition and incubated at 37 °C was included in the experiment for comparison (Fig. 2). Growth curves of *E. coli* cultures under the same temperatures were also recorded and are shown in Fig. S5.

**FIG 2.**
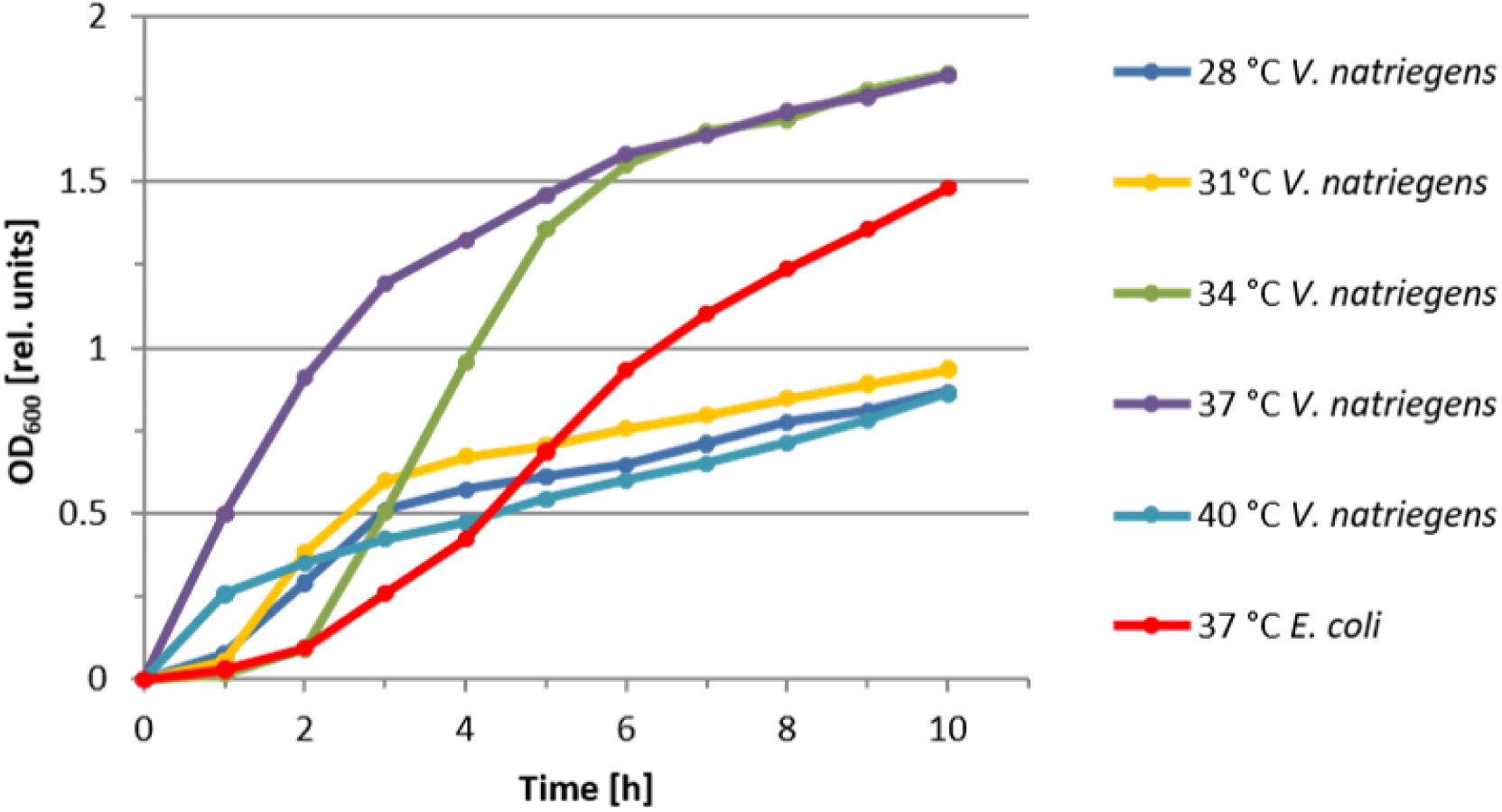
Effect of incubation temperature on the growth dynamics of *V. natriegens* in LB medium supplemented with 2.5 % NaCl. For comparison, the growth curve of *E. coli* grown in LB medium supplemented with 1.0 % NaCl at 37 ° C is shown.

Based on this result we conclude that 37 °C is the optimal temperature to support best growth of *V. natriegens*. In particular, the log-phase is extremely short at this temperature. A similar final OD is reached also at 34 °C, but with a markedly delayed log-phase. 31 °C or 40 °C incubation temperatures significantly inhibits *V. natriegens* growth.

### pH Optimization

Next, we wanted to investigate the pH tolerance and pH optimum for best growth conditions. LB medium cultures supplemented with 2.5 % NaCl and the pH adjusted to 5.0-10.0 in 0.5 steps were inoculated with the same starting amounts of bacteria and incubated at 37 °C. Samples were taken at the indicated times and the OD was measured. *E. coli* grown under identical conditions and with the pH adjusted to 6.0 was included in the experiment for comparison (Fig. 3). Growth curves of *E. coli* cultures under the same pH values were also recorded and are shown in Fig. S6.

**FIG 3.**
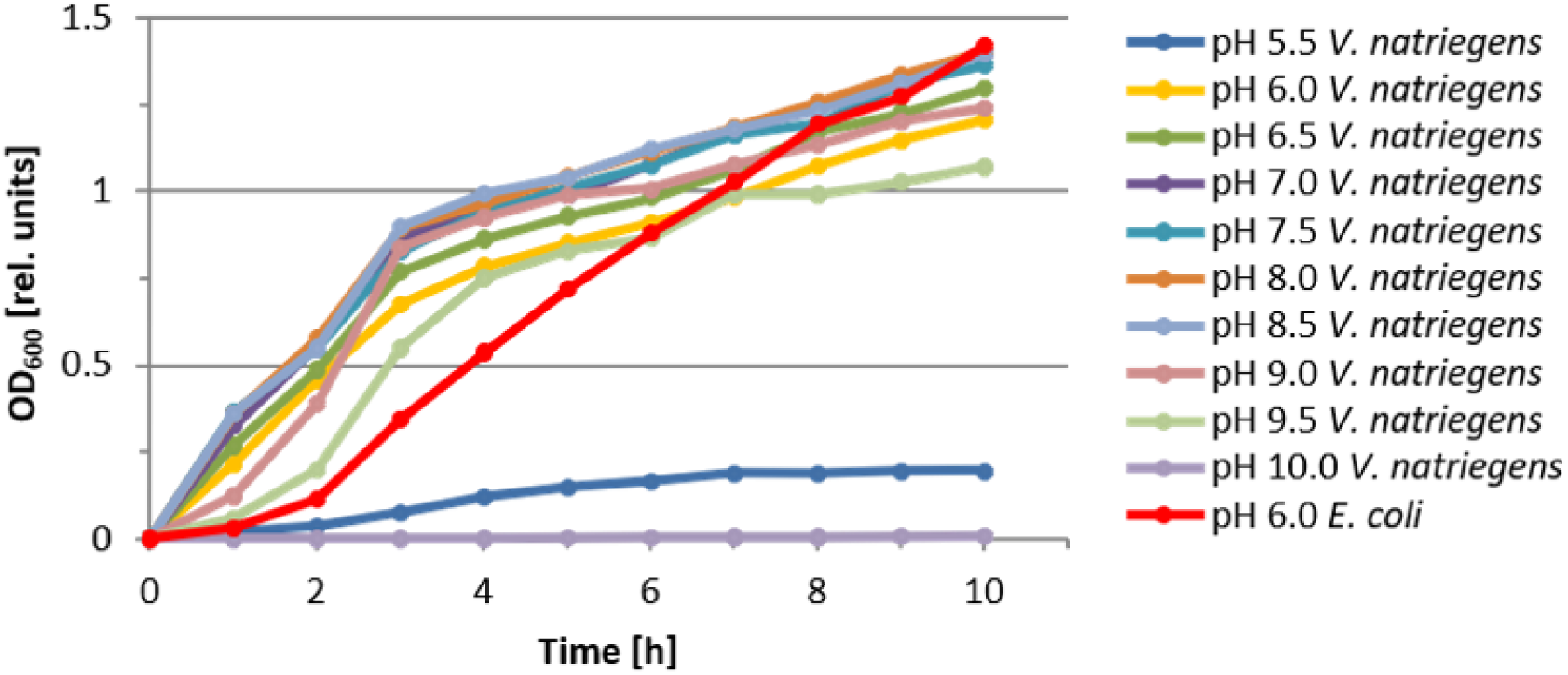
Effect of the pH on the growth dynamics of *V. natriegens* in LB medium supplemented with 2.5 % NaCl. For comparison, the growth curve of *E. coli* grown in LB medium supplemented with 1.0 % NaCl at 37 ° C and pH 6.0 is shown.

With the exception of pH 5.5, all *V. natriegens* cultures grew well, indicating a wide range of pH tolerance. Even at a pH 9.5 (pale green curve), bacterial growth was seen. Only at the extremes (pH 5.5 and pH 10) no (pH 10; grey curve) or only minimal (pH 5.5; dark blue curve) growth was scored.

It can be concluded that *V. natriegens* possesses a broad and efficient pH-regulating metabolism and thus a wide tolerance for different environmentally occurring pH values. For experimental laboratory conditions, using pH 7.0 - 8.5 values are best (violet, turquois, orange, and pale blue in Fig. 3).

### Salt Requirement

*V. natriegens* is known as a halophilic bacterium (1, 2, 3). In order to precisely measure the degree of halophilic behavior we varied the NaCl concentration from 1.0 % to 4.0 % in steps of 0.5 % using LB medium. For comparison we included *E. coli* grown at 1.0 % in this experiment (Fig. 4). Growth curves using NaCl concentrations of 1.5 %, 2.0 %, 2.5 %, 3.0 %, and 4.0 % showed very similar dynamics, with 2.5 % reaching the highest final OD of 1.021. Only the lowest NaCl concentration of 1.0 % resulted in markedly slower growth. *E. coli*, with a NaCl concentration of 1.0 % reached after 10 h and after a long lag phase a similar final OD. *E. coli* growth under identical NaCl concentration, a pH 7.0 and incubation at 37 °C was also recorded (Fig. S7).

**FIG 4.**
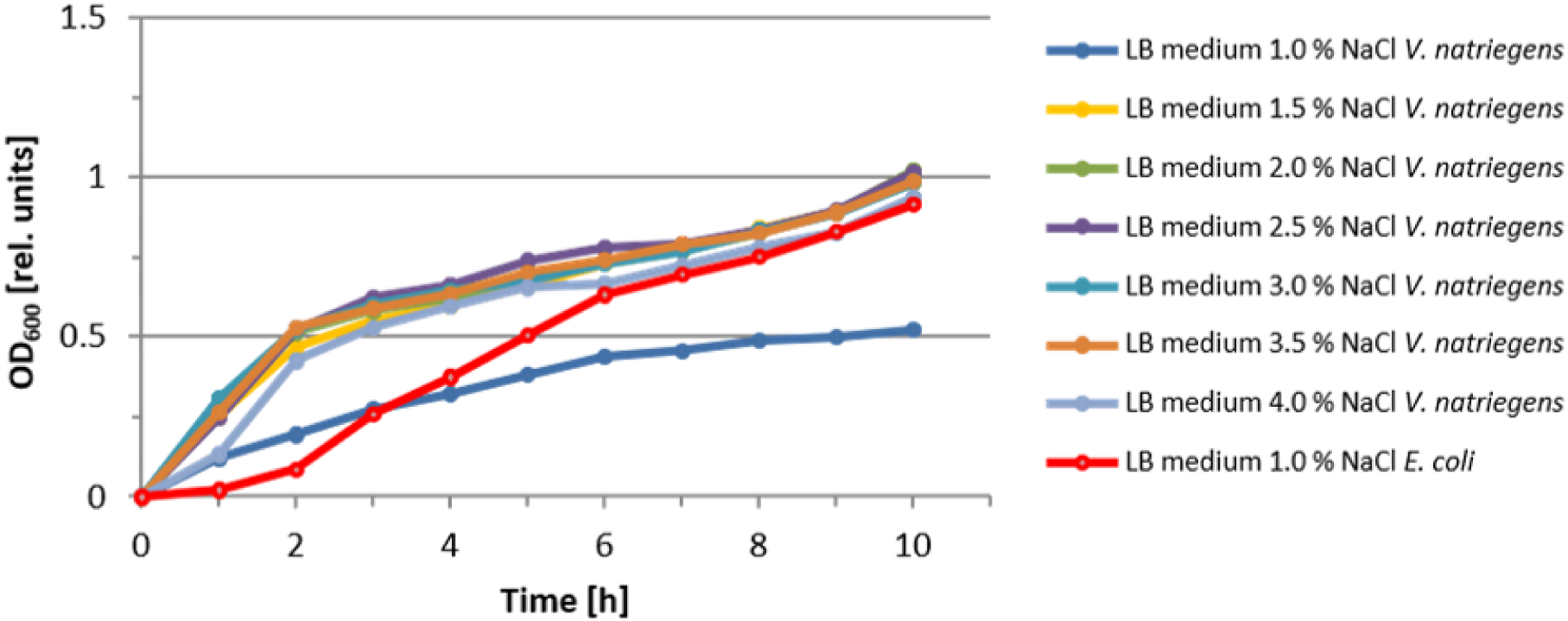
Effect of NaCl concentration on the growth dynamics of *V. natriegens* in LB medium supplemented with the indicated NaCl concentration. For comparison, the growth curve of *E. coli* grown in LB medium supplemented with 1.0 % NaCl at 37 °C and pH 7.0 is shown.)

Based on this result we conclude a NaCl concentration of 2.5 % to be the best for *V. natriegens* growth.

### Microscopy

We wanted to comparatively visualize bacterial morphology of *V. natriegens* and *E. coli* and its dependency on cell density in culture (Fig. 5A). Cells were diluted and optical densities at 600 nm were measured. Cells were analyzed under the light microscope with a 400-fold magnification at the indicated times. As expected, *V. natriegens* cells grew faster than *E. coli* and formed a much denser cell lawn after 2 hours. Next, we wanted to visualize growth behavior of individual cells / cell clusters microscopically. Fig. 5B shows two microphotographs of *V. natriegens* taken of the same region at 0 Minutes and after 30 Minutes. Individual cells and cell clusters can be seen having duplicated within this short time frame (see arrows).

**FIG 5A.**
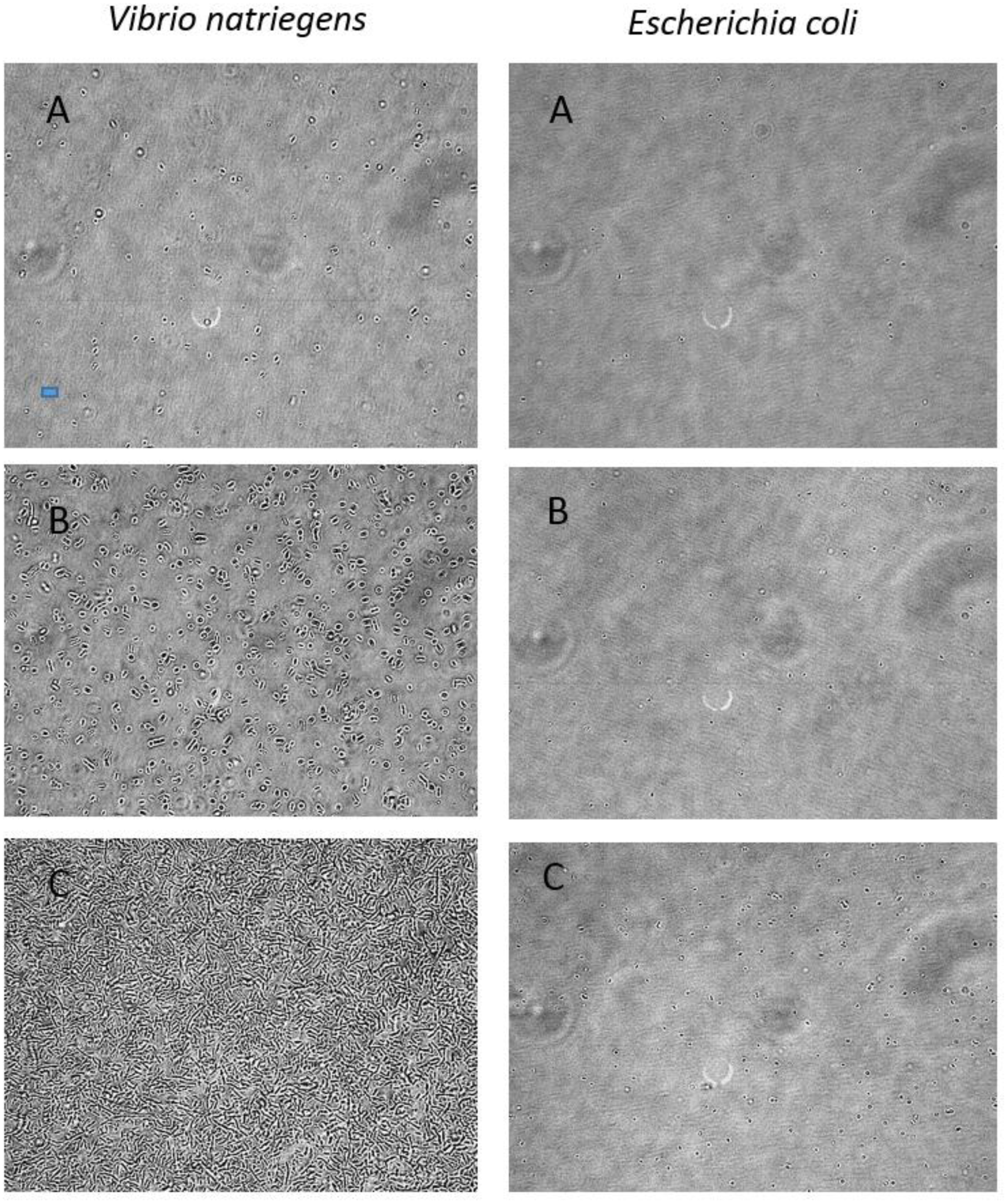
Growth dynamics of *V. natriegens* and *E. coli* in direct comparison. Cells were grown as described in Materials and Methods. (A) Starting dilution at “0 hours” starting dilution (*V. natriegens* OD_600_ = 0.127; *E. coli* OD_600_ = 0.106); (B) Cultures at 1 hour (*V. natriegens* OD_600_ = 0.600; *E. coli* OD_600_ = 0.108); (C) Cultures at 2 hours (*V. natriegens* OD_600_ = 0.404; *E. coli* OD_600_ = 0.108). Blue bar in left photograph A corresponds to 7.5 μm. All photos were taken at the same magnification of 400-fold.

**FIG 5B.**
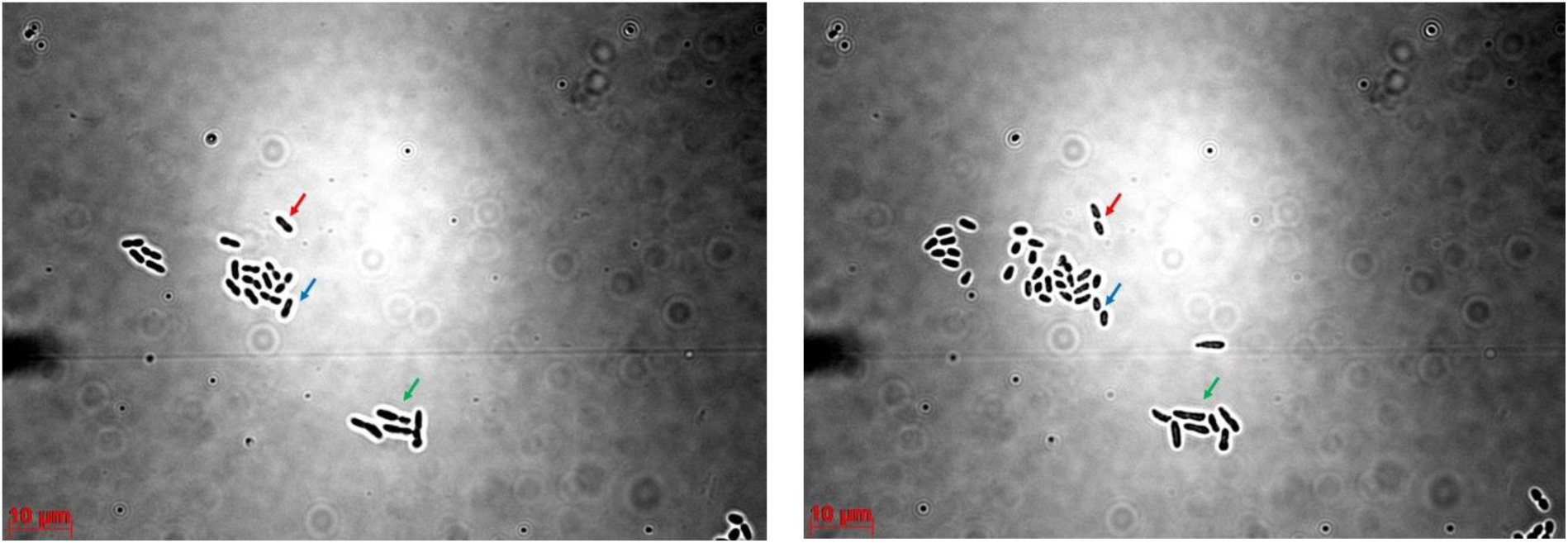
Growth dynamics of *V. natriegens.* The same region was photographed at “0 minutes” (left microphotograph) and at 30 minutes (right microphotograph). Arrows in color indicate corresponding individual dividing cells / cell clusters. Magnification was 1.000-fold. Scale bars in red correspond to 10 μm.

### Cytochrome C Oxidase Activity of V. natriegens

We wanted to study if *V. natriegens* displays cytochrome C oxidase activity. For comparison E. coli, which is known to miss this activity, was included in the study. Fig. 6A shows that *V. natriegens* displays a strong cytochrome C oxidase activity, whereas *E. coli* doesn’t.

**FIG 6A.**
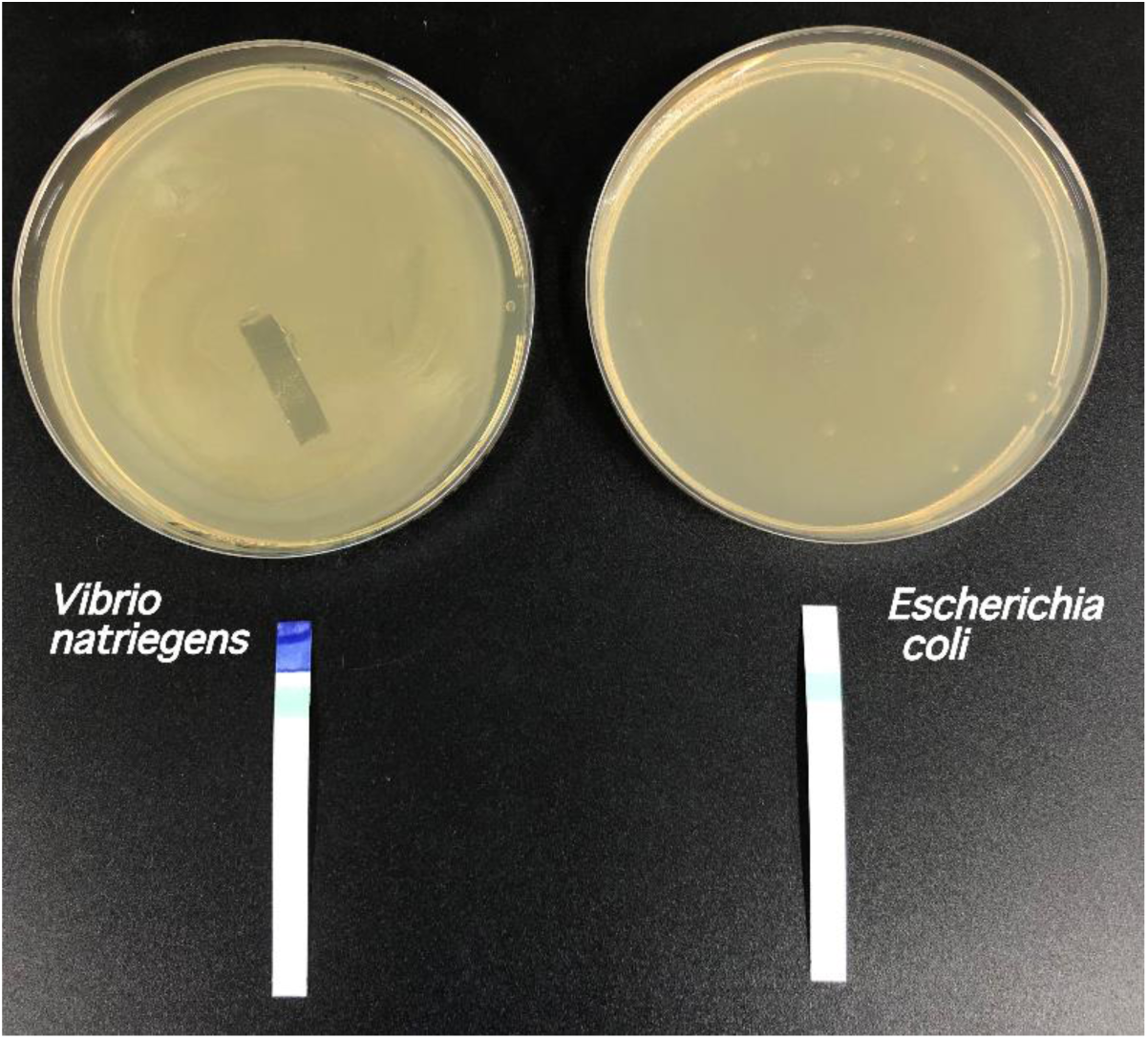
Comparison of cytochrome C oxidase activities of *V. natriegens* and *E. coli.* Petri dishes were incubated for 18 h at 37° C with the indicated bacteria. The *V. natriegens* lawn was thicker when compared with the *E.coli* lawn, which, however, had sufficient amounts of bacteria for carrying the following test: Test strips (Merck Bactident® Oxidase) were placed on the petri dish and incubated for 1 minute. The *V. natriegens* indicates strong cytochrome C oxidase activity, whereas *E. coli* doesn’t. The stripe in turquoise indicates the positive control.

### Missing Catalase Activity of V. natriegens

We wanted to study if *V. natriegens* displays catalase activity. For comparison *E. coli*, which is known to display this activity, was included in the study. Fig. 6B shows that *V. natriegens* displays only a low (if any) catalase activity, whereas *E. coli* shows a strong catalase activity, as expected.

**FIG 6B.**
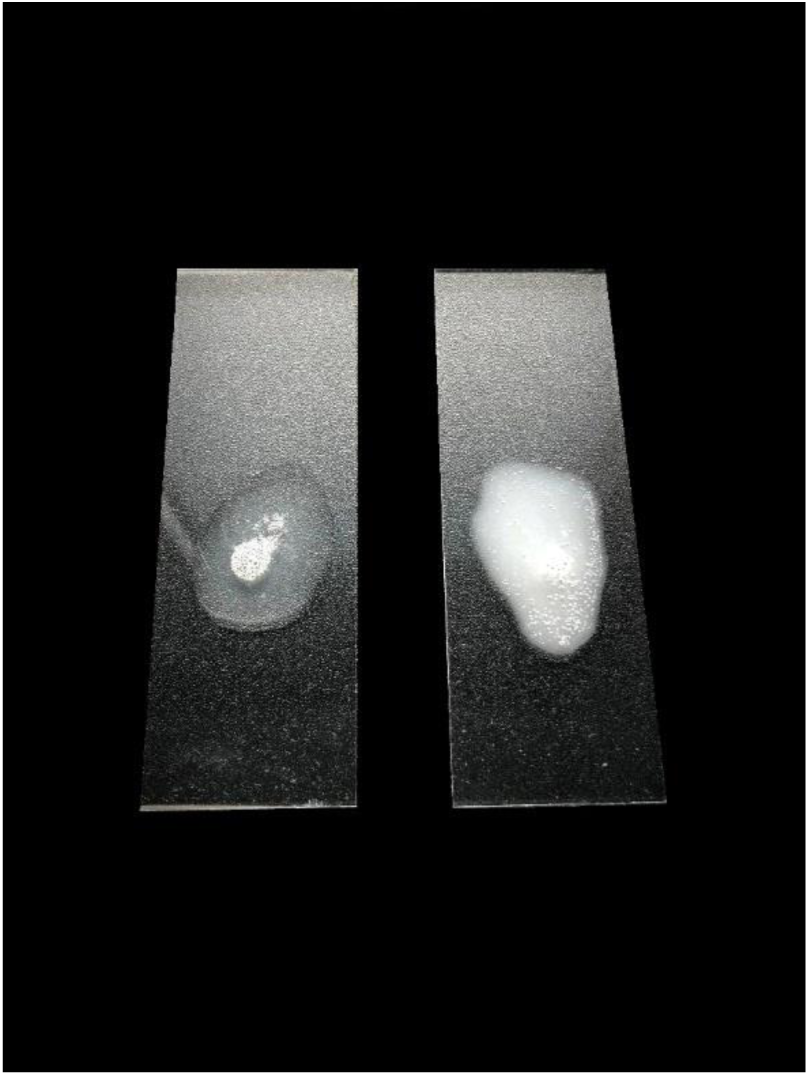
Comparison of catalase activities of *V. natriegens* and *E. coli.* Approximately the same number of bacteria were applied to microscopy slides (*V. natriegens*: left slide; *E. coli*: right slide). Five drops of 3 % hydrogen peroxide were then added and bacteria and hydrogen peroxide were thoroughly mixed with an inoculating loop. Formation of foam is seen for *E.coli*, indicating a high catalase activity. Formation of foam in the case of *V. natriegens* is minimal, indicating only a low catalase activity.

## DISCUSSION

*V. natriegens* is the fasted-growing nonpathogenic bacterium isolated so far (3). Intrigued by extremely fast growth dynamics, *V. natriegens* was proposed and is already applied as a novel host for applications in molecular biology, biotechnology, genomics and industrial applications, for example for the high-level production of recombinant proteins (4, 8, 13). Despite the quest to utilize *V. natriegens* for such useful application, its growth requirements and growth behavior is still poorly understood.

We therefore intended to study systematically and in more detail conditions for optimal growth, with the goal to establish simple and standardized growth conditions applicable in any routine laboratory setting and excluding advanced fermentation technology for extremely high bacterial and protein yield. We analyzed i) various growth media composition, ii) incubation temperature, iii) pH dependence, and iv) salt concentration requirements for optimal growth of *V. natriegens* strain DSMZ 759.

As a result of the studies, the following optimal conditions were established: LB medium with 2.5 % NaCl, pH 7.0 - 8.0 and incubation at 37 °C under aerobic conditions. Incubation temperatures above 37 °C slows growth significantly. Incubation temperatures below 37 °C slows growth, but at a lower rate. Incubation at or below 28 °C should be avoided.

Under such optimized, standard laboratory conditions, a doubling time of t_d_ = 13.6 minutes was observed for *V. natriegens* measured in mid-log growth phase.

The optimized conditions presented here for the growth of *V. natriegens* can be easily applied in any standardly equipped laboratory.

## MATERIALS AND METHODS

### Bacterial Strains

*Vibrio natriegens* Strain DSMZ 759

*E. coli* strain XL1 Blue

### Media and Culture Conditions

*V. natriegens* was stored at −80°C as 50% (vol/vol) glycerol stocks. For cultivation bacteria were streaked out or resuspended, respectively, on different agar plates and in the different media listed below.

LB (Luria-Miller; Carl Roth; X969.2) agar contained in liter^-1^: 10 g tryptone, 5 g yeast extract, 10 g NaCl and 15 g agar. The amount of NaCl (Carl Roth; 9265.1) in liter^-1^ was varied between 10 g, 15 g and 25 g.

LB medium containing in liter^-^1: 10 g tryptone, 5 g yeast extract and 10 g NaCl. The NaCl amount in liter^-1^ was varied between 10 g, 15 g and 25 g.

Nutrient agar contained per liter: 5 g peptone from casein (Sigma; 70172-100G), 3 g yeast extract (Carl Roth; 2363.2) and 15 g agar (Sigma; A4800-500G). The NaCl concentration was varied between 5 g, 15 g and 25 g per liter.

Nutrient medium contained 5 g peptone from casein and 3 g yeast extract per liter. The NaCl amount per liter was varied between 5 g, 15 g and 25 g NaCl.

M9 minimal agar contained per liter: 12.8 g Na2HPO4 (Carl Roth; T876.1), 3 g KH2PO4 (Carl Roth; P018.1), 15 g agar (Sigma; A4800-500G), 0.5 g NaCl (Carl Roth; 9265.1), 1 g NH4Cl (PanReac AppliChem.; 141121.1210), 20 ml of 20 % D(+)-Glucose (Carl Roth; HN06.3), 2 ml of 1 M MgSO4 (PanReac AppliChem; 142486.1211) and 0.1 ml of 1 M CaCl2 (Carl Roth; A119.1).

LB (Luria-Miller; Carl Roth; X969.2) agar contained: 10 g tryptone liter-1, 5 g yeast extract liter-1 and 15 g agar liter-1. The concentration of NaCl (Carl Roth; 9265.1) was varied between 1.0 % (10 g NaCl liter-1), 1.5 % (15 g NaCl liter-1) and 2.5 % (25 g NaCl liter-1). LB medium containing 10 g tryptone liter-1 and 5 g yeast extract liter-1. The NaCl concentration was varied between 1.0 % (10 g NaCl liter-1), 1.5 % (15 g NaCl liter-1) and 2.5 % (25 g NaCl liter-1).

Nutrient agar contained 5 g peptone from casein (Sigma; 70172-100G) liter^-1^, 3 g yeast extract (Carl Roth; 2363.2) liter^-1^ and 15 g agar (Sigma; A4800-500G) liter^-1^. The NaCl concentration was varied between 0.5 % (5 g NaCl liter^-1^), 1.5 % (15 g NaCl liter^-1^) and 2.5 % (25 g NaCl liter^-1^).

Nutrient medium contained 5 g peptone from casein liter^-1^ and 3 g yeast extract liter^-1^. The NaCl concentration was varied between 0.5 % (5 g NaCl liter^-1^), 1.5 % (15 g NaCl liter^-1^) and % (25 g NaCl liter^-1^).

M9 minimal agar contained 12.8 g Na_2_HPO_4_ (Carl Roth; T876.1) liter^-1^, 3 g KH_2_PO_4_ (Carl Roth; P018.1) liter^-1^, 15 g agar (Sigma; A4800-500G) liter^-1^, 0.5 g NaCl (Carl Roth; 9265.1) liter^-1^, 1 g NH_4_Cl (PanReac AppliChem.; 141121.1210) liter^-1^, 20 ml of 20 % C_6_H_12_O_6_ (Carl Roth; HNO6.3), 2 ml of 1 M MgSO_4_ (PanReac AppliChem.; 142486.1211) and 0.1 ml of 1 M CaCl_2_ (Carl Roth; A119.1).

### Light Microscopy

LB medium with a total concentration of 2.5 % NaCl for *V. natriegens* and 1.0 % for *E. coli* were separately inoculated starting with a part of the bacterial lawn from agar plates containing 2.5 % NaCl. After cultivation overnight at 37 ^o^C with shaking at 140 rpm, 50 µL of the cultures were used for inoculation of 20 mL LB medium containing 2.5 % NaCl for *V. natriegens* and 1.0 % NaCl for *E. coli*. The cultures were placed into a Petri dish and over a time period of 10 hours photographs were taken with an inverse microscope (Motic, Wetzlar, Germany). Photographs at time points 0, 1, and 2 hours are shown in Fig. 5A. At these time points, the OD_600_ of the cultures were determined.

Another culture of *V. natriegens* was diluted 1:10,000 and 10 µL of the dilution were put on a diagnostic object slide (Fig 5B). Photographs were taken over a period of 30 min (AXIO Imager D1/Z1; Carl Zeiss Microscopy GmbH; Jena, Germany) with a magnification of 1000-fold and with an oil-immersion objective.

## SUPPLEMENTAL MATERIAL

Supplemental material for this article may be found at https://…………

## ACKNOWLEDGEMENTS

The authors like to thank Gerd Bange and Alex Lepak for providing *V. natriegens* strain DSMZ 759.

This research received no specific grant from any funding agency in the public, commercial, or not-for-profit sectors.

## SUPPLEMENTAL MATERIAL

### Comparison of Growth Dynamics of V. natriegens and E.coli

We wanted to compare the growth dynamics of *V. natriegens* with those of *E. coli* (Fig. S1). *V. natriegens* was grown in LB Medium supplemented with 2.5 % NaCl at pH 7.0 and *E. coli* was grown in LB Medium supplemented with 1.0 % NaCl at pH 7.0. at temperatures of 28 °C, 31 °C, 34 °C, 37 °C and 40 °C, respectively. For each experiment, 1,000 μl of overnight cultures were transferred into 25 mL LB medium with 2,5 % NaCl in shaker flask vials and incubated under shaking at 140 rpm on a reciprocating shaker (MaxQ 6000, Thermo Scientific) under aerobic conditions. At the indicated times, 1,000 μl were transferred to a cuvette and the OD = 600 was measured on the spectrophotometer uniSPEC2, LLG Labware. After measurement, the sample transferred back to its liquid culture and the incubation in the shaker flaks continued.

Fig. S1 shows that for all temperatures tested *V. natriegens* displays faster growth dynamics and higher end-point ODs than *E. coli.* The 40 °C temperature (E) shows the only exception, in that the *V. natriegens* growths dynamics is impaired at a higher degree than that of *E. coli*. For 34 °C and 37 °C temperatures, the end-log phase cell densities are highest, both for *V. natriegens* and for *E. coli.* The highest difference in growth dynamics between *V. natriegens* and *E. coli* are observed at 28 °C (A).

**SUPPLEMENTARY FIG S1.**
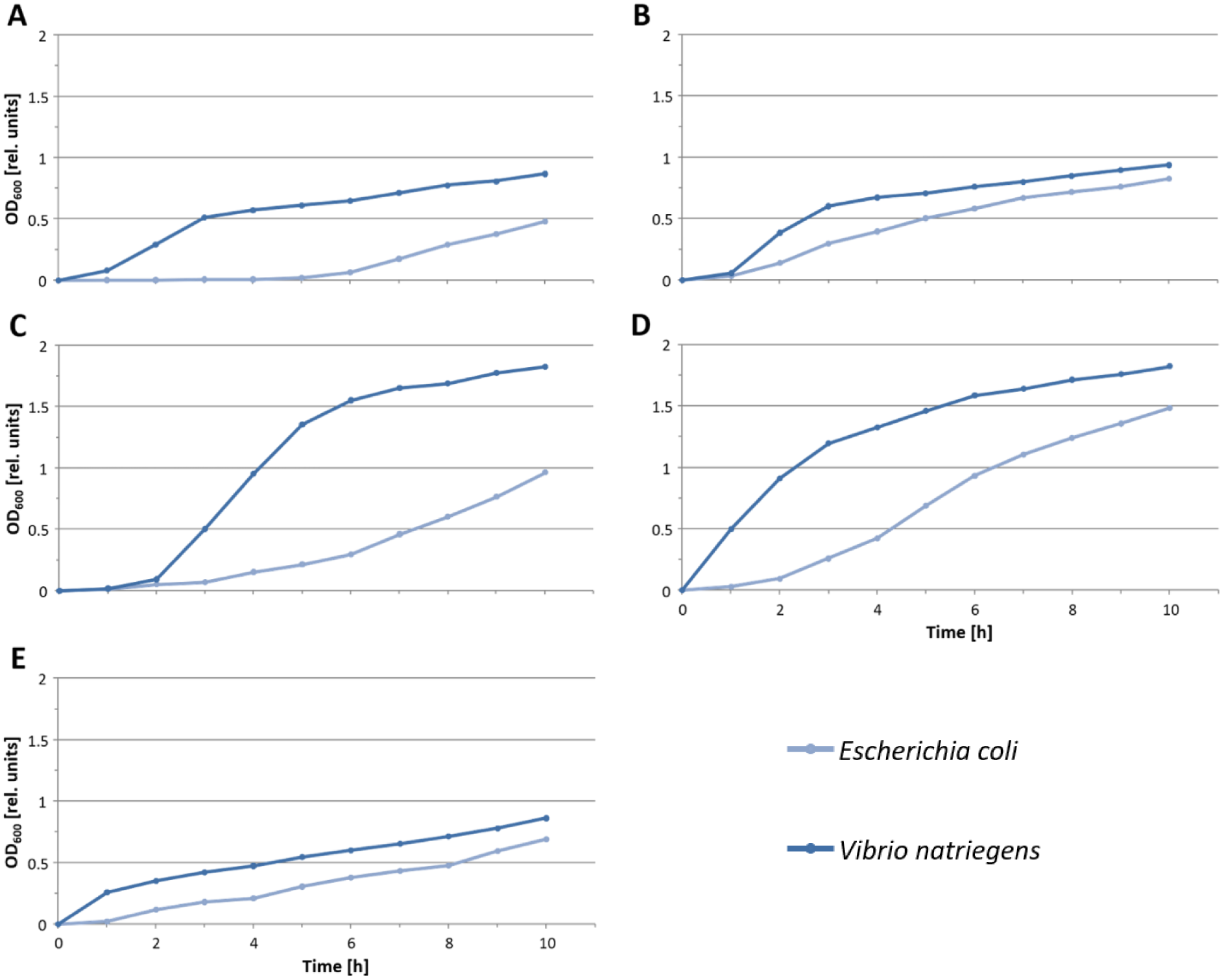
Comparison of growth dynamics of *V. natriegens* and *E. coli* in LB standard medium at pH 7.0. *V. natriegens* was grown with 2.5 % NaCl and *E. coli* was grown with 1.0 % NaCl. (A) 28 °C, (B) 31 °C, (C) 34 °C, (D) 37 °C, (E) 40 °C.

### Comparison of pH on Growth Dynamics of V. natriegens and E.coli

Next, we investigated the effect of the pH on the growth dynamics of *V. natriegens* in direct comparison with those of *E. coli* (Fig. S2). *V. natriegens* was grown in LB Medium supplemented with 2.5 % NaCl and *E. coli* was grown in LB Medium supplemented with 1.0 % NaCl. Incubation temperature for both bacterial species was 37 °C.

For each experiment, 1,000 μl of overnight cultures were transferred into shaker flask vials adjusted to a different pH as indicated in graphs A - I of Fig. S2 and incubated under shaking at 140 rpm on a reciprocating shaker MaxQ 6000, Thermo Scientific under aerobic conditions. At the indicated times, 1,000 μl were transferred to a cuvette and the OD_600_ was measured on the spectrophotometer uniSPEC2, LLG Labware. After measurement, the sample was transferred back to its liquid culture and the incubation in the shaker flaks continued.

For all tested pH, with the exception of pH 5.5 (A) and pH 6.0 (B), the growth of *V. natriegens* outrivals that of *E. coli*. This is an interesting finding indicating that *E. coli* is less prone to acidic environments than *V. natriegens.* On the other hand, *E. coli* growth is markedly impaired on the basic end of the spectrum, starting at pH 9.0 (H) and stronger impaired at pH 9.5 (I) compared with *V. natriegens.* Unsurprisingly, at pH 10.0 none of both bacteria Growth (J).

**SUPPLEMENTARY FIG S2.**
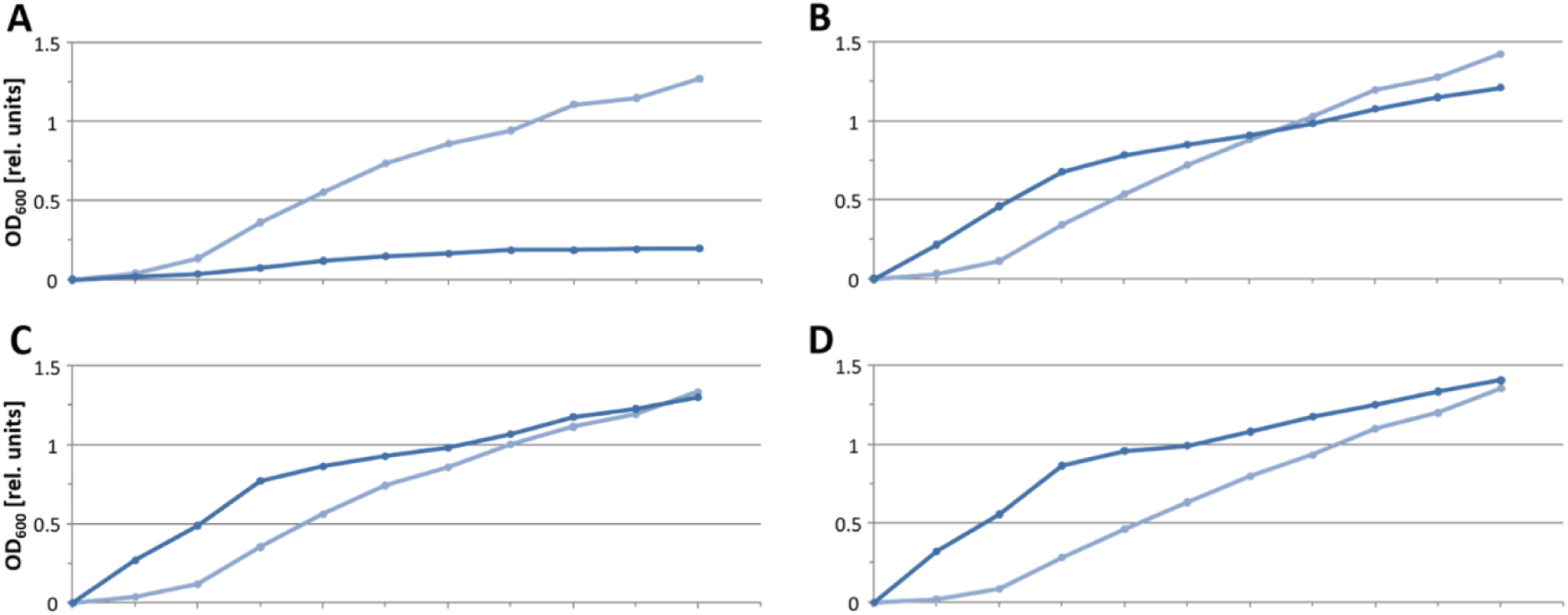
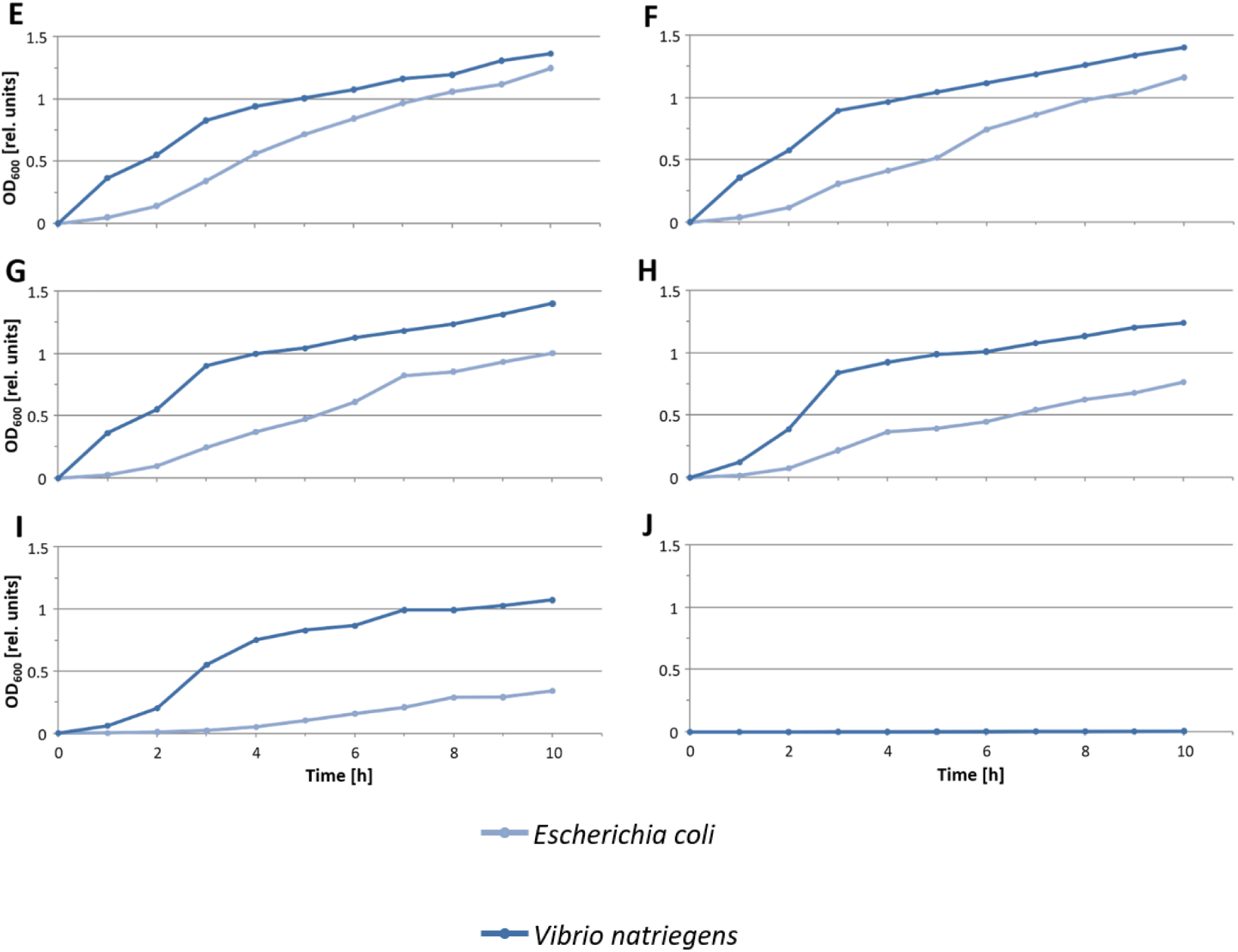
Comparison of growth dynamics of *V. natriegens* and *E. coli* in LB standard medium with varying pH conditions. *V. natriegens* was grown with 2.5 % NaCl and *E. coli* was grown with 1.0 % NaCl incubated at 37 °C. (A) pH 5.5, (B) pH 6.0, (C) pH 6.5, (D) pH 7.0 (E) pH 7.5, (F) pH 8.0, (G) pH 8.5, (H) pH 9.0, (I) pH 9.5, (J) pH 10.0.

### Comparison of Growth Dynamics of V. natriegens and E.coli in Dependency of NaCl

In the next experiment we investigated the salinity requirements of *V. natriegens* in direct comparison with those of *E. coli* (Fig. S3). *V. natriegens* and *E. coli* were grown in LB Medium with varying NaCl concentrations at pH 7.0 and at 37 °C. The LB Medium was supplemented with NaCl to a final concentration of 1.0 %, 1.5 %, 2.0 %, 2.5 %, 3.0 %, and 3.5 % und 4.0 %, respectively.

For each experiment, 1,000 μl of overnight cultures were transferred into 25 ml shaker flask vials and incubated under shaking at 140 rpm on a reciprocal shaker (MaxQ6000, Thermo Fisher) under aerobic conditions. At the indicated times, 1,000 μl were transferred to a cuvette and the OD600 was measured on the spectrophotometer uiSPEC2, LGG Labware. After measurement, the sample was transferred back to its liquid culture and the incubation in the shaker flaks continued.

Only for the lowest NaCl concentration of 1 % was the *E. coli* growth superior to the one of *V. natriegens* (A). For all other NaCl concentrations used, *V. natriegens* overruns *E. coli*’s growth. At a NaCl concentration of 4 %, the growth of *E. coli* is severely inhibited, whereas growth of *V. natriegens* is not impaired (G).

**SUPPLEMENTARY FIG S3.**
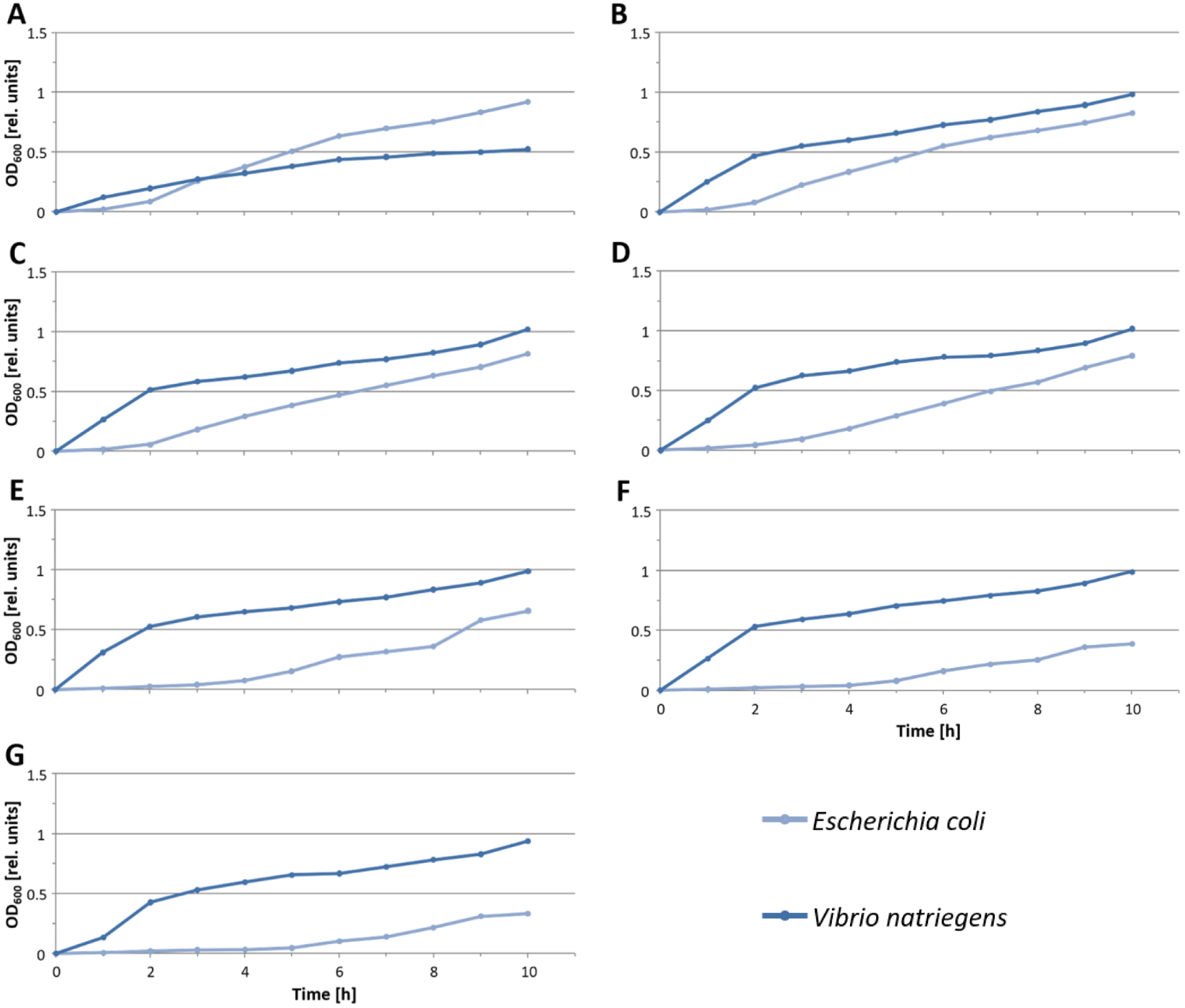
Comparison of growth dynamics of *V. natriegens* and *E. coli* in LB standard medium with varying NaCl concentrations. *V. natriegens* and *E. coli* were grown at 37 °C with a pH 7.0. (A) 1.0 % NaCl, (B) 1.5 % NaCl, (C) 2.0 % NaCl, (D) 2.5 % NaCl, (E) 3.0 % NaCl, (F) 3.5 % NaCl, (G) 4.0 % NaCl.

### E. coli Medium Optimization

Growth curves of *E.coli* were also recorded in order to test three media (LB medium, nutrient medium, M9 minimal medium) each supplemented with 1.0 %, 1.5 %, and 2.5 % NaCl, with the exception of M9 minimal medium. Incubation was at 37 °C and pH was 7.0 (Fig S4). LB medium supplemented with 1.5 % NaCl reached the highest OD_600_. With the exception of M9 minimal medium, all other media with the three salt concentrations can be used for good growth of the *E. coli* bacterial cells.

**SUPPLEMENTARY FIG S4.**
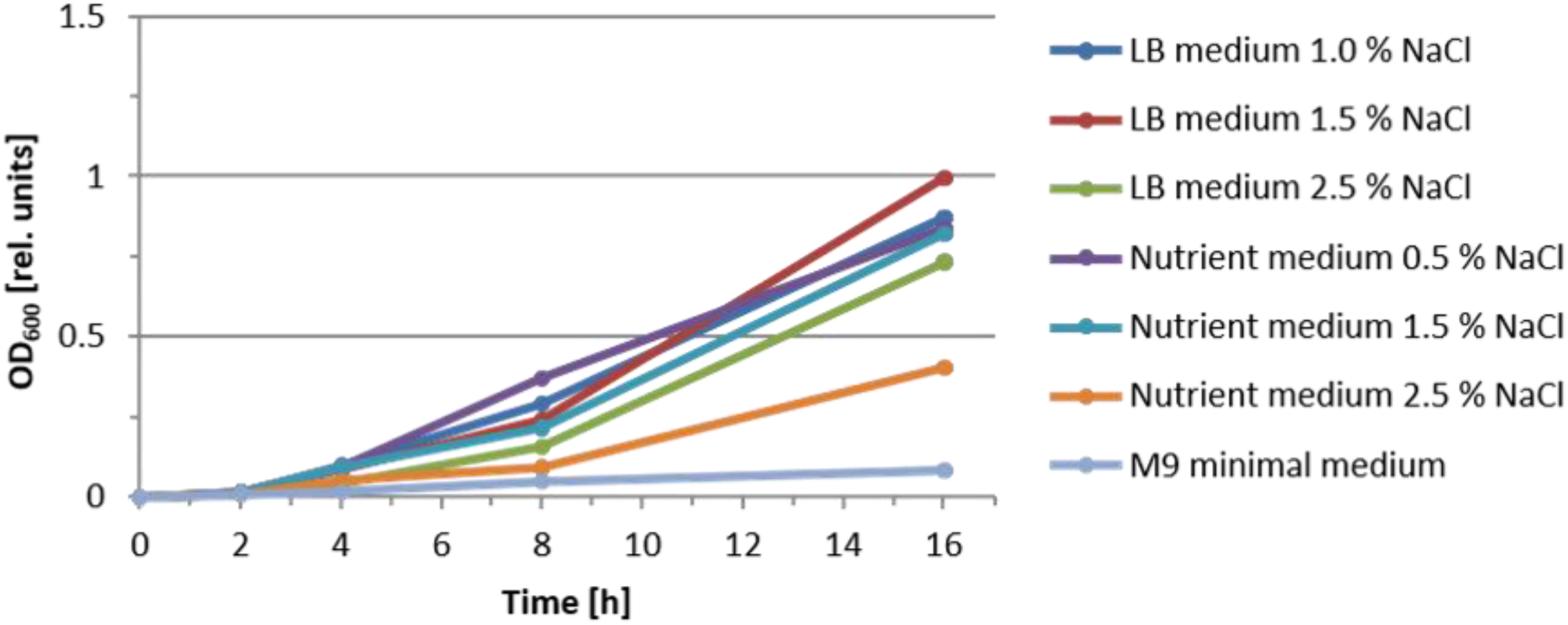
Growth of *E. coli* in different standard media, supplemented with 1.0 %, 1.5 %, and 2.5 % NaCl over time. The M9 minimal medium was not supplemented with NaCl.

### E. coli Growth Temperature Optimization

Growth curves of *E.coli* were also recorded in order to test incubation temperature effects. Incubation was performed at 37 °C and pH was 7.0 (Fig. S5). Best growth conditions were seen at 37 °C, where after 10 hours an OD_600_ of 1.5 was reached. Also, growth dynamics in the mid-log phase was superior to the other temperatures tested. At 40 °C, significant loss of cell growth was observed.

**SUPPLEMENTARY FIG S5.**
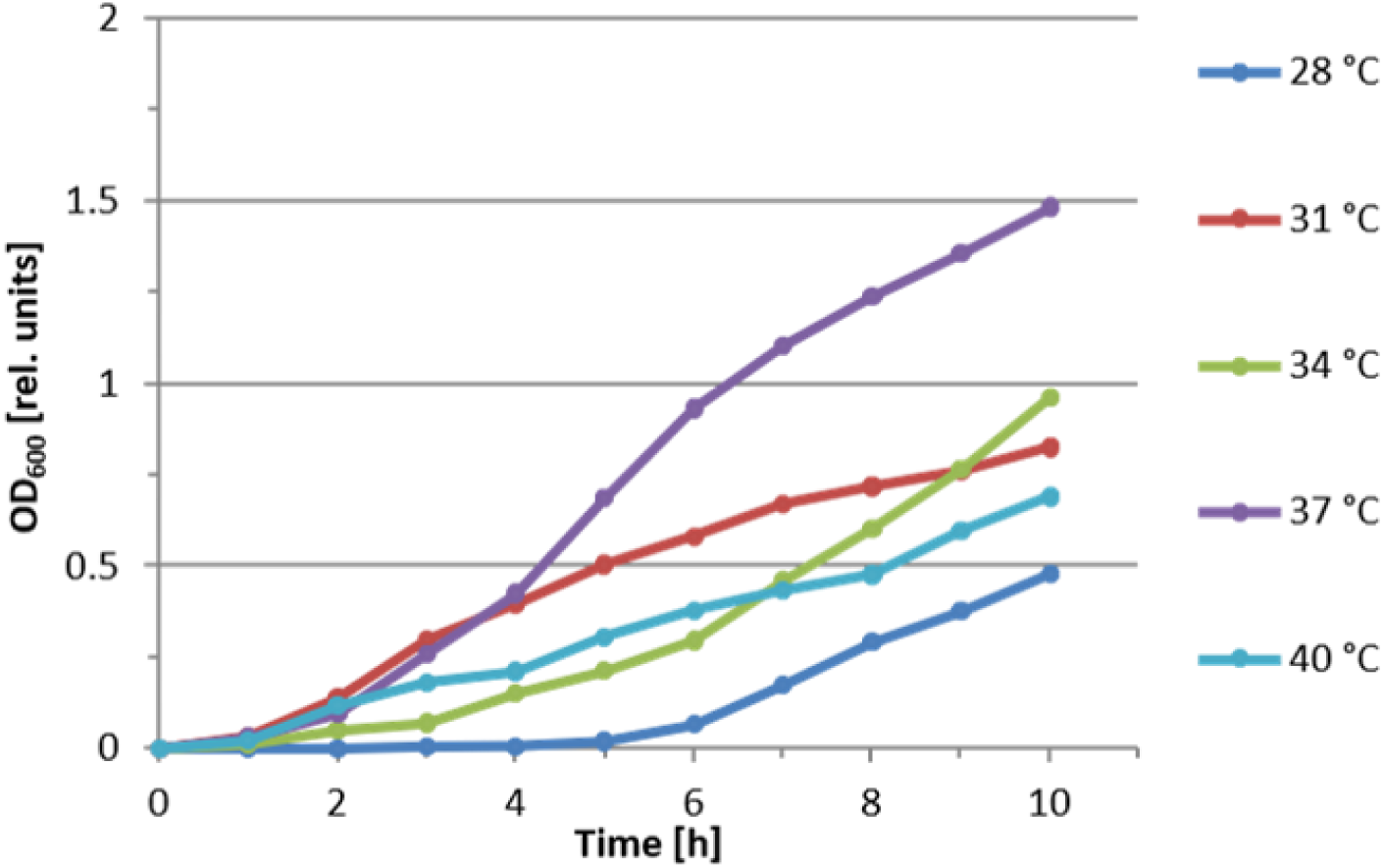
Effect of incubation temperature on the growth dynamics of *E. coli* in LB medium supplemented with 1.0 % NaCl and incubated at pH 7.0.

### E. coli Growth pH Optimization

Growth curves of *E.coli* were also recorded in order to test the growth implication of different pH used. Cultivation took place in LB medium supplemented with 1.0 % NaCl at 37 °C (Fig. S6). The result shows that *E. coli* grows well in a wide range of pH, from pH 5.5 to pH 8.0. Best growth was recorded at pH of 6.0. It can be concluded, that the pH over this wide range is ideal to promote the growth of *E. coli*. At a pH of 9.5, growths is markedly inhibited and at a pH of 10.0, no growth can be observed.

**SUPPLEMENTARY FIG S6.**
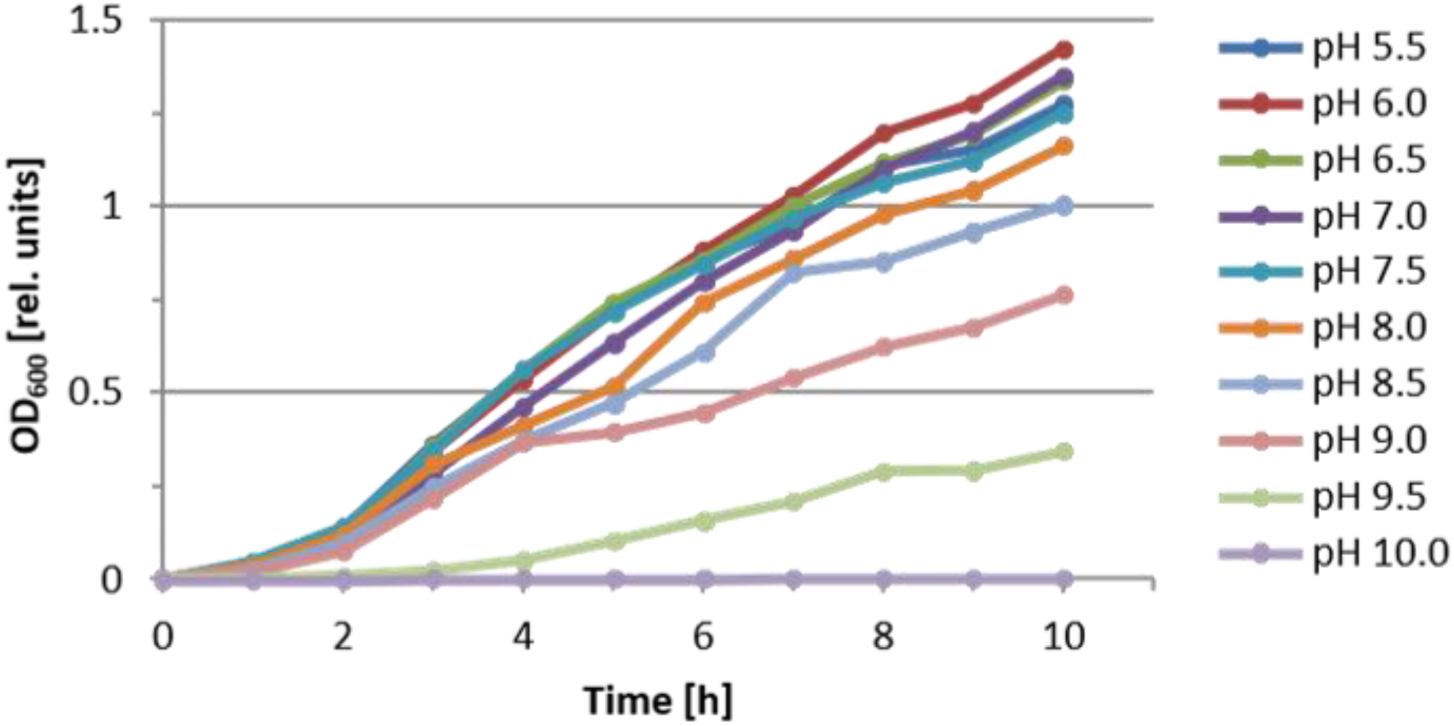
Effect of the pH on the growth dynamics of *E. coli* in LB medium supplemented with 1.0 % NaCl at 37 °C.

### E. coli Growth NaCl Concentration Optimization

Growth curves of *E.coli* were also recorded in order to test the implication of different salt concentrations on the growth dynamics of bacterial cells. Cultivation was performed in LB medium supplemented with the indicated NaCl concentration at 37 °C and a pH of 7.0 (Fig. S7). The salt concentration has a stronger effect on *E. coli* growth than on *V. natriegens* growth.

The optimal NaCl concentration for E.coli growth was 1.0 %, with descending growth dynamics from 1.5 % to 4.0 %. Concentrations of 3.5 % and 4.0 %, respectively, showed severely impaired growth curves.

**SUPPLEMENTARY FIG S7.**
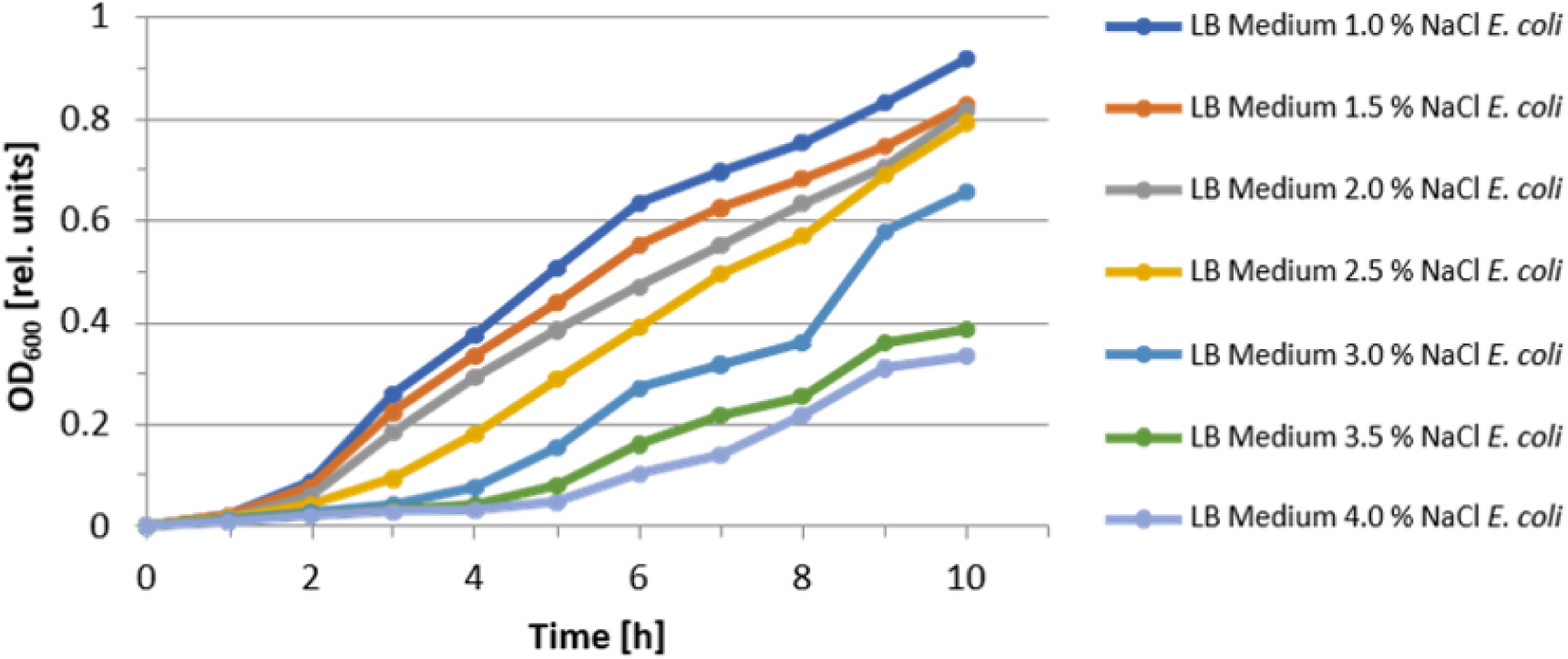
Effect of NaCl concentration on the growth dynamics of *E. coli* in LB medium supplemented with the indicated NaCl concentration. Cultures were incubated at 37 °C at pH 7.0.

